# Cognitive abilities are associated with rapid dynamics of electrophysiological connectome states

**DOI:** 10.1101/2024.01.15.575736

**Authors:** Suhnyoung Jun, Stephen M. Malone, William G. Iacono, Jeremy Harper, Sylia Wilson, Sepideh Sadaghiani

## Abstract

Time-varying changes in whole-brain connectivity patterns, or connectome state dynamics, hold significant implications for cognition. However, connectome dynamics at fast (> 1Hz) timescales highly relevant to cognition are poorly understood due to the dominance of inherently slow fMRI in connectome studies. Here, we investigated the behavioral significance of rapid electrophysiological connectome dynamics using source-localized EEG connectomes during resting-state (N=926, 473 females). We focused on dynamic connectome features pertinent to individual differences, specifically those with established heritability: Fractional Occupancy (i.e., the overall duration spent in each recurrent connectome state) in beta and gamma bands, and Transition Probability (i.e., the frequency of state switches) in theta, alpha, beta, and gamma bands. Canonical correlation analysis found a significant relationship between the heritable phenotypes of sub-second connectome dynamics and cognition. Specifically, principal components of Transition Probabilities in alpha (followed by theta and gamma bands) and a cognitive factor representing visuospatial processing (followed by verbal and auditory working memory) most notably contributed to the relationship. We conclude that the specific order in which rapid connectome states are sequenced shapes individuals’ cognitive abilities and traits. Such sub-second connectome dynamics may inform about behavioral function and dysfunction and serve as endophenotypes for cognitive abilities.

## 1. Introduction

Cognitive processes are inherently dynamic, with a substantial portion of these dynamics unfolding at speeds of few 100 milliseconds or faster. It stands to reason that ongoing or spontaneous neural processes constituting most brain activity, including large-scale connectome dynamics, affect cognition (Raichle and Gusnard, 2005). Indeed, great functional significance of large-scale network dynamics for cognitive performance (Sadaghiani et al., 2015; Kucyi et al., 2018) and inter-individual differences therein (Eichenbaum et al., 2020; Jun et al., 2022) has been established using fMRI. In fact, investigations of functional connectome dynamics and their functional significance have been dominated by fMRI due to its exceptional spatial resolution. Unfortunately, the temporal dynamics captured by the fMRI-derived slow and indirect measure of neural activity, or BOLD signal, misses the rich temporal dynamics that occur on cognitively more relevant sub-second timescales.

These time-varying dynamics in large-scale functional connectivity can be characterized as flexible changes in connectome states, representing varying strength of connectivity between specific sets of brain regions within the whole-brain connectome, which occur repeatedly over time. Recently, data-driven approaches, such as Hidden Markov Modeling (HMM), have been employed to identify temporally recurrent connectome states with state-specific mean and covariance from the observed time series of different brain regions.

Prior fMRI connectome studies have established that a wide range of behaviors and cognitive processes are linked to such recurrent state dynamics, encompassing both the temporal organization of connectome state transitions (Vidaurre et al., 2017; Eichenbaum et al., 2020; Jun et al., 2022) and changes in the spatial organization of connectivity patterns (Sadaghiani et al., 2015; Shine et al., 2016). Particular temporal features of fMRI-derived connectome dynamics, specifically the proportion of the total recording time spent in each connectome state (Fractional Occupancy) and the probability to transition between specific pairs of connectome states (Transition Probability), have not only been linked to behavioral performance (Vidaurre et al., 2017; Eichenbaum et al., 2020; Jun et al., 2022), but also found to be heritable (Vidaurre et al., 2017; Jun et al., 2022). More specifically, our previous fMRI work has established substantial genetic effects (*h*^*2*^ ∼ 40%) and behavioral relevance of Fractional Occupancy and Transition Probability (Jun et al., 2022). We further found preliminary evidence for specific genetic polymorphisms predictive of fMRI-derived Fractional Occupancy and Transition Probability via the regulatory impact of modulatory neurotransmitter systems (Jun et al., *Under Review;* https://doi.org/10.17605/OSF.IO/VF2ZW). This fMRI-based body of literature establishes that *infra-slow* connectome dynamics are endophenotypes driving individually specific cognitive abilities, which in turn are heritable (Han and Adolphs, 2020; Jun et al., 2022). Yet, little is known about the functional significance of *rapid* connectome dynamics.

Non-invasive, real-time methods, i.e., EEG and MEG, allow capturing rapid electrophysiological signals with real-time fidelity. Recently, a growing body of work has established the capability to investigate the spatially informative functional connectome and, importantly, its rapid dynamics in source-space with measures addressing source leakage (static/time-averaged connectome: (de Pasquale et al., 2010; Brookes et al., 2011; Deligianni et al., 2014; Hipp and Siegel, 2015; Wirsich et al., 2017), connectome dynamics: (Baker et al., 2014; Brookes et al., 2014; Sitnikova et al., 2020; Wirsich et al., 2020; Coquelet et al., 2022), for review, see (Sadaghiani and Wirsich, 2020)). Importantly, we have recently shown that Fractional Occupancy and Transition Probability of rapid connectome dynamics derived from source-space EEG are also under significant genetic influence (Jun et al., 2024; Submitted). This subject-specificity of temporal phenotypes of rapid connectome dynamics lead us to question their functional implications for individual differences in cognitive abilities.

In the current study, we focused on the *heritable* time-varying phenotypes of sub-second, electrophysiological connectome dynamics. Heritable features inherently reflect individual differences and further allow us to narrow the particularly large feature space of rich, electrophysiological connectome dynamics that comprise numerous frequency bands. We investigated whether such band-specific dynamic phenotypes contribute to inter-individual variability in cognitive task measures. Specifically, we applied Hidden Markov Modelling (HMM) to obtain discrete brain states using source-reconstructed resting-state EEG data from two cohorts of twins from the Minnesota Twin Family Study (Iacono et al., 1999; Keyes et al., 2009; Wilson et al., 2019). For each canonical frequency band, we focused on previously identified heritable temporal phenotypes (i.e., Fractional Occupancy and Transition Probability; (Jun et al., 2024; Submitted)) and examined their associations with cognitive task measures using canonical correlation analysis. To the best of our knowledge, this is the first source-reconstructed EEG study investigating the behavioral significance of rapid temporal connectome dynamics.

## 2. Materials and Methods

Figure 1 is a schematic representation of the overall approach and analysis subsections.

**Figure 1.**
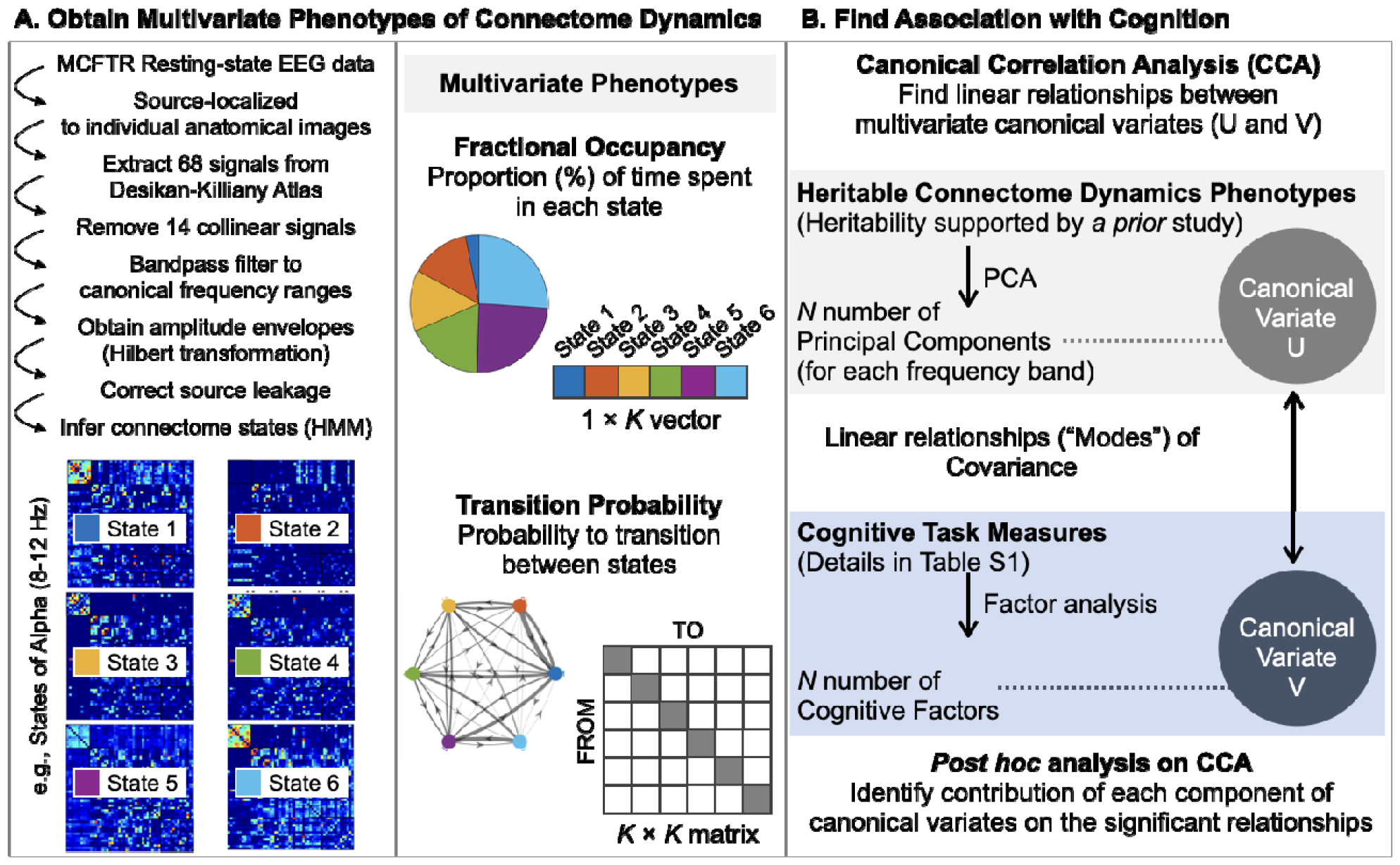
An overview of the analysis pipeline. [A] We used resting-state EEG signals in source-space and structural images sourced from the Minnesota Twin Family Study dataset. Employing a hidden Markov model (HMM), we extracted six discrete connectome states for amplitude timeseries of each canonical frequency band. These states were associated with a state time course for each subject, indicating the likelihood of each state being active. Connectivity matrices reflecting amplitude coupling are shown for each state. These six states are color-coded (blue, red, yellow, green, purple, and light blue) to visually depict their contribution to the connectome dynamics of interest. From this analysis, we obtained two temporal features of rapid connectome dynamics for each frequency band, namely the proportion of time spent in each connectome state (Fractional Occupancy) and the probability matrix describing transitions between all possible pairs of discrete states (Transition Probability). These features were constructed in a multivariate manner to comprehensively represent all states. For further analysis, we retained Fractional Occupancy and Transition Probability in bands in which they were found to be heritable (Jun et al., 2024; Submitted). [B] In order to identify linear relationships (“modes”) between these multivariate dynamic connectome phenotypes and cognitive task measures, we conducted canonical correlation analysis (CCA). This analysis was carried out on dimensionality-reduced data, encompassing frequency-specific *N* principal components of multivariate dynamic connectome phenotypes and *N* cognitive factors.

### 2.1. Subjects and cognitive measures

Participants for the present investigation are from the two independent cohorts of twins from the Minnesota Twin and Family Research (MCTFR) (Iacono et al., 1999; Keyes et al., 2009; Wilson et al., 2019). Twins in both cohorts have been followed periodically since approximately the age of 11. As part of their most recent assessment, participants underwent structural MRI scans in addition to resting EEG recordings. At time of initial recruitment and at each follow up, participants gave written informed consent or assent, if under the age of 18 for their participation.

A total of 1164 subjects in two independent cohorts of MCTFR twins completing virtually identical assessments, 928 had usable and complete EEG data which was source-localized successfully (see **2.3 EEG signal pre-processing and source localization**), thus permitting HMM-based estimation of discrete connectome states. The included subjects (473 females) were 23–40 years of age at time of data acquisition. Subsequently, 463 sex-matched pairs (926 subjects) were formed for heritability analysis as follows: 206 monozygotic (MZ) twin pairs, 112 sex-matched dizygotic (DZ) twin pairs, and 145 pairs of sex-matched unrelated individuals. Each subject entered only one pair. Note that the unrelated individuals are twins whose co-twin lacked complete data and was therefore not part of the analytic sample.

We included 15 summary measures from 8 cognitive tasks completed as part of a whole-day assessment at the MCTFR. Tasks assessed the following domains: verbal and nonverbal intelligence; visual-spatial processing speed; auditory-verbal learning and memory; auditory-verbal attention and memory; visual-spatial attention and memory; visual-spatial learning and memory; response inhibition; and decision-making under uncertain conditions or risk (see *Table S1* for more detailed description for each measure and task). The measures were transformed into z-scores. All 926 subjects had more than 50% of cognitive measures (859 subjects with complete dataset).

### 2.2 MRI and EEG acquisition

Structural MRI data were collected on either 3T Siemens Trio or Prisma MRI scanner (32-channel array head coil) at the Center for Magnetic Resonance Research, University of Minnesota. Three-dimensional T1-weighted sagittal plane anatomical images were acquired using the following magnetization-prepared rapid gradient echo sequence: TR = 2530 ms; TE = 3.65 ms; flip angle = 7°; matrix size = 256 × 256; FOV = 256 mm; GRAPPA = 2; 240 coronal slices with 1-mm isotropic voxels; single shot; interleaved acquisition.

While recording EEG, participants rested comfortably in a darkened room, with their head and neck supported while hearing 55-dB white noise played through headphones. They were instructed to keep their eyes closed and relax. A recorded voice subsequently instructed them to open the eyes or close them at 1-min intervals. A total of 6 min of EEG was collected, 3 min with eyes open and 3 with eyes closed. EEG data were acquired from 61 scalp electrodes arranged according to the International 10/10 system using a BioSemi ActiveTwo system (BioSemi, Amsterdam, The Netherlands) at 1024 Hz. ActiveTwo amplifiers are DC coupled. ActiveTwo signals are monopolar. They were low-pass filtered using a digital 5th-order Bessel antialiasing sinc filter with a cutoff frequency (3-dB attenuation) of 205 Hz. Pairs of electrodes placed above and below the right eye or on the outer canthus of each eye allowed for detecting blinks and other eye movements. Additional electrodes were placed on left and right earlobes, and the average of these signals was derived offline to serve as a reference.

### 2.3. EEG signal pre-processing and source localization

Raw resting-state EEG signals were pre-processed using a monitored automated pipeline (https://www.github.com/sjburwell/eeg_commander) of Minnesota Twin Family Study group and EEGLAB (Delorme and Makeig, 2004) in MATLAB (version R2021b, Mathworks, Inc.). Signals were down-sampled to 256 Hz, filtered with a 0.1 Hz high-pass filter (*firfilt* EEGLAB plugin; 1,286 Kaiser window), and referenced to averaged earlobe signals. A monitored automated pipeline detected four kinds of signal anomalies: disconnected channels/flat signals, interelectrode electrolyte bridging (Tenke and Kayser, 2001), large amplitude deviations, and muscle/cap shift (motion) noise. Descriptives (e.g., temporal variance) were calculated for each electrode and 1-s time range. Data that exceeded four normalized median absolute deviations from the median (Rousseeuw and Croux, 1993) in 25% of a 1s time range or 75% of a given electrode were removed. Among others, this approach is effective in removing periods with head motion artifacts. Ocular correction was performed using independent components (IC) analysis (infomax algorithm; Bell and Sejnowski, 1995) and joint consideration of temporal and spatial signal characteristics. The IC time series and inverse weights were compared with the time courses of the bipolar vertical or horizontal EOG and the inverse weight of a stereotypical blink or horizontal saccade to correct for vertical and horizontal ocular artifacts, respectively. If the squared joint temporal and spatial correlations for an IC exceeded an empirically calculated threshold based on Expectation Maximization (Mognon et al., 2011), that IC was subtracted from the data.

For source localization, we imported preprocessed EEG recordings and MR-based anatomical images into Brainstorm software (Tadel et al., 2011). The EEG signals were resampled to 250 Hz, corrected for DC offsets, linearly detrended, and low-pass filtered at 70 Hz. We manually marked fiducial points, including the anterior commissure (AC), posterior commissure (PC), inter-hemispheric point, nasion (NAS), and left and right pre-auricular points (LPA and RPA), of all subjects using their individual anatomical images to aid coregistration of electrode positions and T1 images. The coregistration was refined by manually moving the electrode positions onto the electrode artifacts visible in the T1 image. We then used the OpenMEEG software (Gramfort et al., 2010) with a symmetric boundary element method (BEM) to calculate a forward model of the skull based on the individual T1 image of each subject (Tadel et al., 2019). Then, we used the Tikhonov-regularized minimum-norm estimation (MNE) inverse method to compute the sources, with default parameter settings for regularization and source depth weighting (Tikhonov parameter = 10%, assumed SNR = 3.0, constrained sources normal to cortex, depth weighting 0.5/max amount 10) (Baillet et al., 2001; Tadel et al., 2019).

### 2.4. Parcellation and Source-leakage correction

We used the Desikan-Killiany Atlas (Desikan et al., 2006) in Brainstrom to average source signals within each of the atlas’ 68 anatomically distinct brain regions. To aid network-level interpretation, we also determined each region’s membership within the seven canonical Intrinsic Connectivity Networks or ICNs (Yeo et al., 2011) based on spatial overlap.

To mitigate source-leakage confounds caused by the blurring of point dipole sources and the spreading of signals across neighboring regions, we excluded regions whose signals were collinear with others based on a QR decomposition, which is commonly used to model the correlation structure of a set of variables, using the *qr* function in Matlab. As a result, 14 regions were excluded from the investigation (*Figure S1*). The remaining 54 regional signals underwent detrending and bandpass filtering within canonical frequency ranges: delta (1-3 Hz), theta (4-7 Hz), alpha (8-12 Hz), beta (13-25 Hz), and gamma (30-45 Hz). Then, we used a symmetric orthogonalization procedure (Colclough et al., 2015) to remove all shared signal at zero lag between the regions. This multivariate method extends previous orthogonalization methods (Brookes et al., 2012; Hipp et al., 2012) and identifies orthogonal time-courses that maintain the closest similarity to the original, unmodified time-series. Finally, amplitude envelopes for each canonical frequency band and brain region were computed using the Hilbert transform, which were then downsampled to 40 Hz (Baker et al., 2014; Hunyadi et al., 2019).

### 2.5. Hidden Markov Modelling of connectome states

The hidden Markov model (HMM) assumes that time series data can be represented by a finite sequence of hidden states. Each HMM-inferred connectome state, along with its corresponding time series, represents a unique connectivity pattern that re-occurs over time. Using the HMM-MAR toolbox (Vidaurre et al., 2016), we applied the HMM to the region-wise EEG amplitude timeseries separately for each frequency band and obtained six discrete connectome states (*K* = 6). While HMMs require an *a priori* selection of the number of states, *K*, the objective is not to establish a ‘correct’ number of states but to strike a balance between model complexity and model fit and to identify a number that describes the dataset at a useful granularity (Quinn et al., 2018). Our previous fMRI-based investigation into connectome heritability (Jun et al., 2022) reported results for two different *K* values (to ensure that outcomes are not limited to a single chosen parameter), namely *K* of 4 and *K* of 6. This choice was in turn informed by prior fMRI literature (Vidaurre et al., 2016; Karapanagiotidis et al., 2020). The choice of *K =* 4 and 6 falls with the range applied in prior HMM studies of EEG and MEG data, which have used *K*s between 3 and 16 (Baker et al., 2014; Vidaurre et al., 2016; Quinn et al., 2018; Hunyadi et al., 2019; Coquelet et al., 2022), where two of the studies used *K* of 6. Therefore, based on the success of our prior fMRI study in revealing heritability within 6-state and 4-state models (Jun et al., 2022), the current study reports results from *K* of 6 (main text) and *K* of 4 (supplementary).

### 2.6. Null model of Hidden Markov Models

To demonstrate that the dynamic trajectory of connectome state transitions is not occurring by chance, we employed a null model. This involved generating 50 simulated state time courses for each frequency band, which were of the same length as the original empirical state time courses. While preserving the static covariance structure, the temporal ordering of states was intentionally disrupted (Vidaurre et al., 2016). It is worth noting that selecting 50 simulations for each of the frequency bands in this analysis represents a rigorous choice in comparison to previous studies (e.g., four simulations in (Vidaurre et al., 2017)). We performed HMM inference with *K*s of 6 (and 4 for replication) on each of these simulated time courses, allowing us to recalculate all above-described temporal and spatial connectome phenotypes at both the group and subject levels. Through this process, we confirmed that the original dataset’s non-random distribution of phenotypes over states represented veridical dynamics as it was absent in the simulated data (see Supplementary *Figure S2* in (Jun et al., 2024; Submitted)).

### 2.7. Multivariate temporal features of the dynamic connectome

The HMM-derived estimates provide a comprehensive set of multivariate temporal features that simultaneously characterized all states of the dynamic connectome. These estimates describe the temporal aspects of connectome dynamics by characterizing the sequence of connectome states, namely the trajectory of the connectome through state space. For each subject, we calculated the Fractional Occupancy (the proportion of total time spent in a given state; 1 × *K*) and Transition Probability (the probability matrix of transitioning between all possible pairs of discrete states; *K* × *K*). While Transition Probability and Fractional Occupancy are not fully independent measures, they contain non-overlapping information about connectome dynamics. For example, a state with particularly high Fractional Occupancy is likely to have high values as initial state and target state in the Transition Probability matrix. Despite such dependence, however, two hypothetical subjects with highly comparable Fractional Occupancy values across the *k* states may still have substantially different sequencing, and thus transition probabilities, across the states. Notably, our previous work demonstrated strong genetic effects specifically on these two temporal phenotypes in fMRI-derived slow functional connectome dynamics (Jun et al., 2022) and EEG-derived rapid connectome dynamics (Jun et al., 2024; Submitted).

### 2.8. Association between dynamic trajectories of states and cognitive performance

Our previous work (Jun et al., 2024; Submitted) has established the substantial and robust heritability of Transition Probability in the theta, alpha, and gamma bands, as well as Fractional Occupancy in the beta and gamma bands. In the present study, we investigated behavioral associations of these heritable connectome dynamics phenotypes using canonical correlation analysis (CCA). CCA tests for linear relationships (or “mode”) between two sets of variables (Hotelling, 1936), and as such as been previously applied to the static connectome and HMM-derived state transitions (Smith et al., 2015; Vidaurre et al., 2017; Eichenbaum et al., 2020; Jun et al., 2022). Here, we trained CCA on the dimensionality-reduced temporal phenotypes of source-space EEG connectome dynamics (U canonical variate matrix detailed below) and cognitive measures (V canonical variate matrix detailed below).

To build the U matrix, we integrated heritable phenotypes of rapid connectome dynamics from all frequency bands. Specifically, these phenotypes encompassed (1) off-diagonals of the *K* × *K* Transition Probability matrix (30 dimension) from the theta and alpha bands, (2) 1 × *K* Fractional Occupancy (6 dimensions) from the beta band, and (3) a composite of off-diagonals of the *K* × *K* Transition Probability and 1 × *K* Fractional Occupancy (36 dimensions) from the gamma band. All variables were normalized (z-scored) before dimensionality-reduction (Wang et al., 2020). For each band separately, Principal Component Analysis (PCA) was applied on the above-described set of multivariate variables, and only principal components (PCs) with eigenvalues exceeding 1 were retained. Subsequently, the U matrix was defined as the PCs aggregated across all bands. Detailed information of PCA on connectome phenotypes for each frequency band can be found in *Figure S2*.

The V matrix encompassed 15 performance measures from 8 cognitive tasks provided by the Minnesota Twin Family Study (see *Table S1* for details of cognitive task measures). Again, measures were z-scored before applying dimensionality-reduction. We adopted dimensionality-reduction methods used for cognitive measures in prior research (Han and Adolphs, 2020; Jun et al., 2022). Specifically, we first conducted the PCA on the cognitive measures to identify the number of PCs with eigenvalues surpassing 1 (*Figure S3A*). Subsequently, we applied the maximum likelihood method for factor analysis, retaining the number of factors, determined based on the preceding PCA. Notably, due to the lack of evidence for the orthogonality among the cognitive measures (as depicted in Figure 2B), we used Promax oblique rotation to adjust them. To further enhance the reliability of our analysis, we calculated factor scores using both ridge regression and Bartlett methods and found that both methods yielded comparable factor scores (*Figure S3B*). Consequently, the V canonical variate comprised of the factor scores derived from the regression method and entered the subsequent CCA.

**Figure 2.**
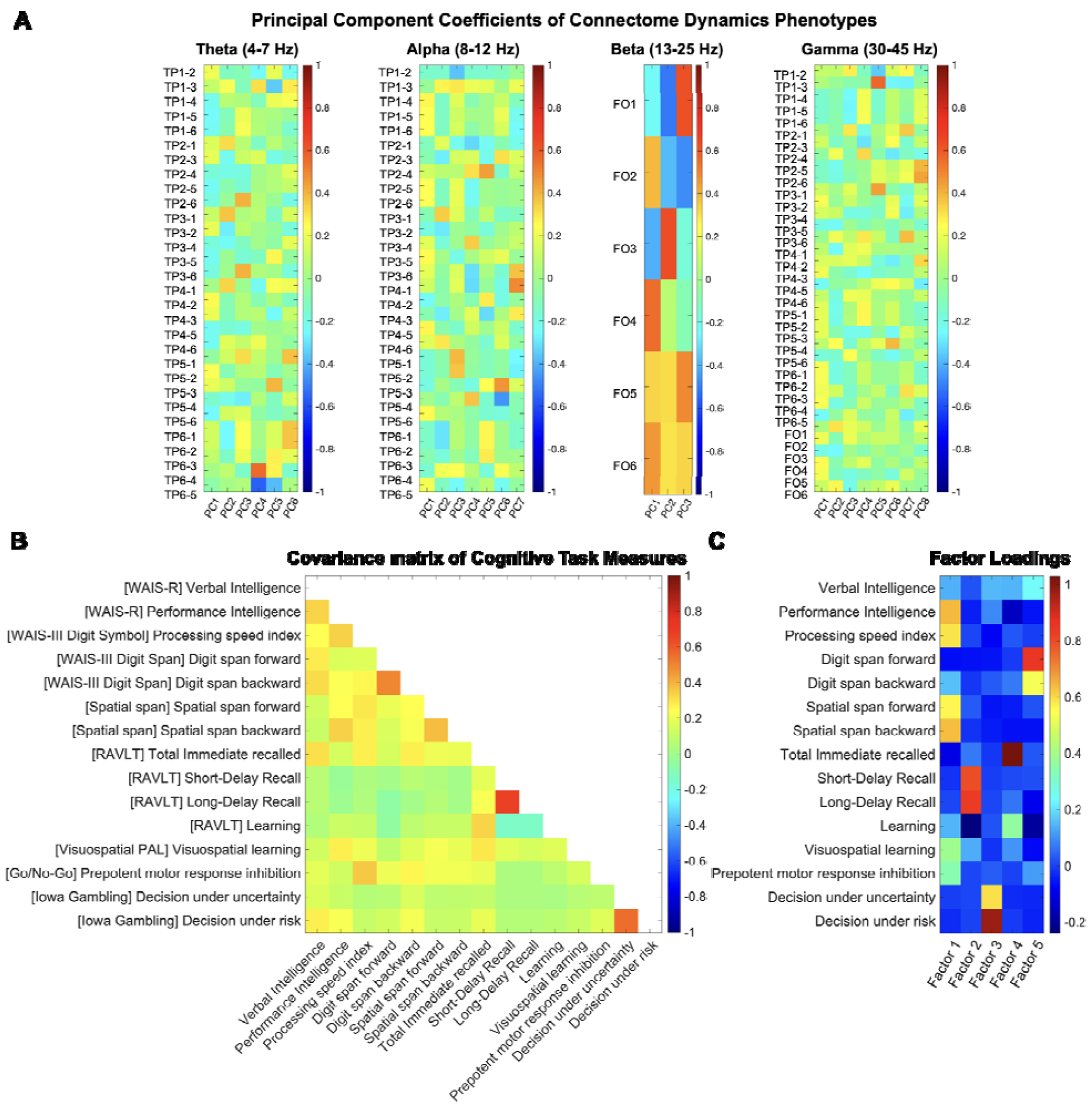
Overview of the dimensionality reduced canonical variates. (A) The principal coefficients matrix displays the weight that each component of the temporal phenotypes has on each of the select number of principal components (eigenvalue > 1). (B) Empirical correlation matrix for the 15 cognitive performance measures from Minnesota Twin Family Study (sample size *N* = 926), color-coded for Pearson’s correlation coefficient. See Table S1 for details of cognitive measures and tasks. (C) The factor loading matrix displays the weight that each cognitive measure has on each of the Factors. FO; Fractional Occupancy, TP; Transition Probability (for example, TP1-2 denotes TP from state 1 to state 2), PC; Principal Component.

To assess the statistical significance of the identified modes of covariation, 10,000 permutations of the rows of U relative to V were performed, while maintaining the within-participant structure of the data. Then, the CCA mode was recalculated for each permutation to generate a distribution of random canonical variate pair correlation values, and each mode of covariation identified in the real data was compared to the random distribution (Smith et al., 2015) (*Figure S4*). Furthermore, post-hoc correlations between the modes and the cognitive factors allowed for determining the contribution of each factor to the given mode and providing further insights into the relationships between the cognitive measures and dynamic connectome features.

## 3. Results

### 3.1. Connectome phenotypes and cognitive measures entering the CCA

In our prior investigation in the same cohort (Jun et al., 2024; Submitted), we established heritability of *temporal* phenotypes of rapid connectome dynamics, specifically Transition Probability in theta, alpha, and gamma bands, as well as Fractional Occupancy in beta and gamma bands. Building upon these findings, we investigated the association between the heritable connectome dynamics phenotypes and cognitive measures using canonical correlation analysis (CCA). Prior to conducting CCA, we applied PCA to the phenotypes of each band separately. This resulted in 6 principal components (PCs) for the theta band (accounting for 81.24% of total variance), 7 for the alpha band (accounting for 86.79% of total variance), 3 for the beta band (accounting for 87.36% of total variance), and 8 for the gamma band (accounting for 88.14% of total variance). These 24 PCs were aggregated to build a canonical variate for CCA. Figure 2A provides a comprehensive illustration of the specific contributions of connectome phenotypes to each PC. Further information of PCA on connectome phenotypes for each frequency band can be found in *Figure S2*.

As for the cognitive measures, we applied dimensionality-reduction (factor analysis following PCA) and retained five cognitive factors. Figure 2C provides insights into the extent to which each cognitive measure contributed to these factors. With cautious acknowledgment that labeling reduced dimensions is suboptimal by nature, we characterize the factors as follows: Factor 1: “Visuospatial Processing” (indicated by high positive loadings across several tasks sharing visuospatial demands); Factor 2: “Verbal Memory” (indicated by high positive loadings of Short and Long Delay Recall measures from Rey Auditory Verbal Learning Task (RAVLT)); Factor 3: “Reward-based decision making” (indicated by high positive loadings of Iowa Gambling Task measures); Factor 4: “Verbal working memory” (indicated by high positive loading of Total Immediate Recall measure from RAVLT); and Factor 5: “Auditory working memory” (indicated by high positive loadings on WAIS-III Digit Span task measures).

### 3.2. Temporal phenotypes of EEG connectome dynamics are associated with cognition

The CCA analysis conducted on the 24 PCs of connectome phenotypes and 5 cognitive factors revealed a significant linear association, commonly referred to as a “mode,” after adjusting for age and sex. The canonical coefficient (*r)* of the mode was found to be .25 with a corresponding *p* value of .015 (Figure 3A). The identified mode underwent non-parametric statistical significance testing (10,000 permutations) establishing significance at *p* = .0012 (*Figure S4*).

**Figure 3.**
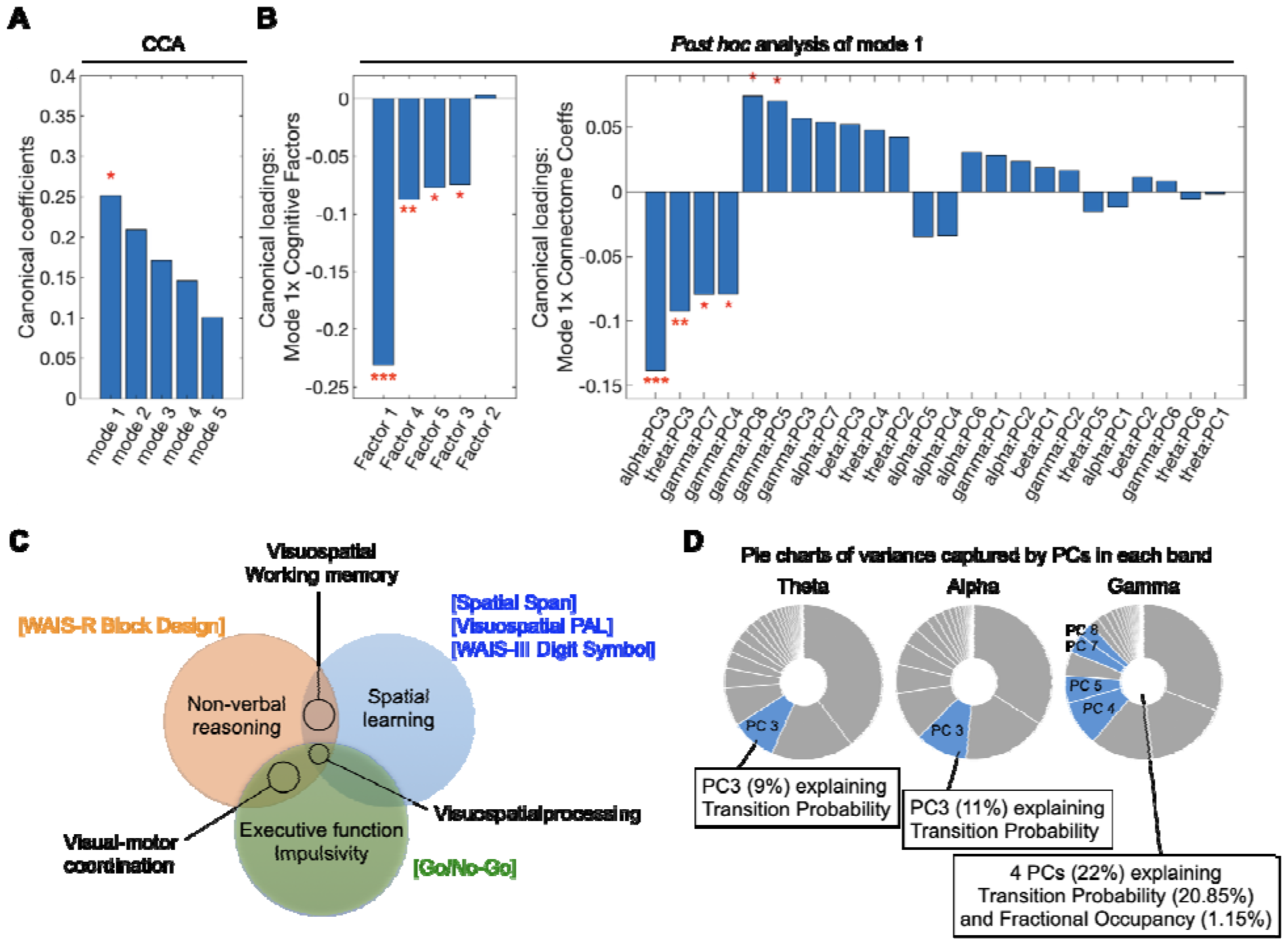
Heritable temporal dynamic connectome phenotypes are related to cognition. Canonical correlation analysis (CCA) finds the maximum linear correlation between two multi-dimensional canonical variates: U and V, adjusting for age and sex. Canonical variate U is defined as 24 principal components (PCs) of connectome dynamics phenotypes across all frequency bands. Canonical variate V is defined as five cognitive factor loadings, each representing different cognitive domain. (A) CCA identified one significant mode of association (B) The significant mode had contributions from four cognitive Factors, namely Factor 1: “Visuospatial Processing”, Factor 3: “Reward-based decision making”, Factor 4: “Verbal working memory”, and Factor 5: “Auditory working memory”. On the connectome side, contributions to the significant mode came from multiple PCs spanning alpha, theta, and gamma bands (C) The Venn diagram illustrate the relationships between different sets of cognitive task measures constructing Factor 1, which contributed significantly to the canonical mode (cf. Figure 2C). Each circle represents cognitive domains measured by the cognitive task(s) of the same color (tasks are labeled in square brackets). The overlapping areas of the circles represent the cognitive domain that is common to multiple cognitive tasks. (D) The pie charts visualize the contribution of band-specific PCs to the canonical mode. Here, each complete pie comprises the total variance in the heritable features of the given frequency band (cf. Figure 2A), and the blue areas reflect the proportion of this variance that significantly contributed to the canonical mode. * *p* < .05, ** *p* < .01, *** *p* < .005.

Subsequent *post hoc* correlation analyses were performed to identify differential contributions (“canonical loadings”) of PCs and cognitive factors to the mode (Figure 3B). We found that Factor 1 (“Visuospatial Processing”; canonical loadings (*r*) = -.23, *p* = 1.11e-12) contributed the most to the significant CCA mode, followed by Factor 4 (“Verbal working memory”; *r* = -.09, *p* = .007), Factor 5 (“Auditory working memory”, *r* = -.08, *p* = .020), and Factor 3 (“Reward-based decision making”; *r* = -.07, *p* = .024). Further, we found significant contributions of PCs from theta, alpha, and gamma bands: alpha PC3 (*r* = -.14, *p* = 2.40e-05), theta PC3 (*r* = -.09, *p* = .005), gamma PC7 (*r* = -.08, *p* = .015), gamma PC4 (*r* = -.08, *p* = .016), gamma PC8 (*r* = .07, *p* = .024), and gamma PC5 (*r* = .07, *p* = .032).

## Discussion

Our current interest in the link between fast, transient connectome dynamics at sub-second timescales and cognition arises from the potential of these timescales to facilitate cognitive functioning and provide endophenotypes of translational relevance. Prior fMRI work has shown that slow time-varying connectome phenotypes may serve as promising endophenotypes for cognitive abilities (Vidaurre et al., 2017; Eichenbaum et al., 2020; Jun et al., 2022). Moving beyond infra-slow timescales, our recent work has established substantial genetic influence on the phenotypes of rapid EEG connectome dynamics (Jun et al., 2024; Submitted). In the present study, we investigated the relationship between these heritable phenotypes – characterizing overall occurrence and sequencing of rapid connectome states– and a broad range of cognitive task measures. Our findings suggest that electrophysiological connectome state transitions unfolding at multiple rapid speeds collectively contribute to shape cognitive abilities.

One of the methods that investigates large-scale brain dynamics with direct electrophysiological techniques has been microstates, denoting recurrent, spatially diffuse sensor-level topographies that transition rapidly (every ∼40-200 ms) (Lehmann et al., 2009; Coquelet et al., 2022). These EEG microstates were found to be predictive of interindividual variability in cognitive abilities (Muthukrishnan et al., 2016; Kim et al., 2021) and linked to neurodegenerative and psychiatric disorders (Hatz et al., 2015; da Cruz et al., 2020). Yet, the diffuse topographies reflected in microstates fall short of informing about connectome states that encompass spatially localized coactivations across networks of brain regions. However, recent advances in methods have significantly improved the study of rapid connectome dynamics at cognitively highly relevant timescales (Baker et al., 2014; Brookes et al., 2014; Sitnikova et al., 2020; Wirsich et al., 2020; Coquelet et al., 2022). Despite these methodological advances, the role of rapid source-space connectome dynamics in individuals’ cognitive abilities has remained elusive.

Such a role in cognitive abilities is likely, given the observation that rapid connectome dynamics are genetically determined to a similar degree than infra-slow connectome dynamics (source-space EEG: Fractional Occupancy in beta (44%) and gamma (40%) bands and Transition Probability in theta (38%), alpha (63%), and gamma (40%) bands (Jun et al., 2024; Submitted) compared to fMRI: Fractional Occupancy (*h*^*2*^=39%) and Transition Probability (*h*^*2*^=43%). (Jun et al., 2022)). Confirming this predicted cognitive significance, our CCA findings unveiled that the heritable phenotypes across multiple frequency bands are significantly associated with various cognitive factors. This finding not only showcases the ability of source-space EEG to capture the functional significance of rich sub-second temporal dynamics, but also demonstrates that rapid connectome transitions occurring at different speeds (i.e., theta, alpha, and gamma bands) *collectively* contribute to this relationship.

Importantly, our data suggest that Transition Probability holds greater functional significance than Fractional Occupancy; along with the third PC of alpha and theta bands that represent the differential contributions of Transition Probability elements, the four gamma-band PCs that contributed to the significant mode of association captured largely variance in Transition Probabilities (Figure 3D). This is likely due to the fact that Transition Probability captures the temporal transition patterns, i.e., the specific sequencing of states, that depict the dynamic changes in interactions between brain regions. These dynamic patterns likely underlie a diverse functional repertoire in the brain, which is relevant for a wide range of behavioral and cognitive outcomes (Deco et al., 2013; Petersen and Sporns, 2015).

The most substantial contribution to the connectome-cognition association emerged from the “Visuospatial Processing” factor (*r* = -.23) and PC3 of the alpha band (*r* = -.14). This observation may be attributed to the extensive role of the alpha rhythm in shaping perception and cognition through the modulation of cortical excitability and subsequent signal processing of neural populations. In particular, alpha oscillation power is known to impact performance in tasks involving visuospatial processing, attention, working memory, and other higher order cognitive control functions (Palva and Palva, 2011; Klimesch, 2012; Mathewson et al., 2012; Sadaghiani and Kleinschmidt, 2016). Alternatively, or additionally, this observation may arise because visuospatial processing is heavily represented among the neurocognitive measures included in our study. As depicted in Figure 2C, the “Visuospatial Processing” factor contains high positive loadings from most of the measures (5 out of 8 tasks) and encompasses various other cognitive functions, including visuospatial working memory, visual-motor coordination, and other higher order cognitive domains (Figure 3C). Moreover, the substantial heritability effect size of the alpha band phenotype (63%) relative to other connectome dynamics phenotypes (38∼44%) (Jun et al., 2024; Submitted) may further influence this finding.

Interestingly, the effect size of this behavioral association with rapid connectome dynamics (*r* = .25, *p* = .015) is similar to that with slow connectome dynamics (*r* = .23, *p* = 5.41e-11; (Jun et al., 2022)). While direct comparisons between these findings are not feasible due to different populations and distinct sets of cognitive measures included in each study, it is interesting to note that the “Language” factor contributed the most (*r* = -.21) to the relationship with slow connectome dynamics, whereas “Visuospatial Processing” factor contributed the most (*r* = -.23) to the relationship with rapid connectome dynamics (Figure 3B).

Our study is subject to several limitations and methodological considerations. While we defined connectome features separately for canonical frequency bands, this approach does not assume or necessitate the bands to be discretely separable or oscillatory in nature. The approach is equally compatible with the view that the bands represent electrophysiological processes at different speed within a larger 1/f spectrum. Further, we defined the boundaries of the frequency bands according to common conventions in the field rather than according to the individual subjects’ power spectrum. While defining the bands individually may strengthen the observed associations to cognition, the non-individualized approach should not invalidate the current findings. Other considerations concern the choice of connectome features and cognitive measures. While our study incorporated all cognitive task measures available from MCTFR, the cognitive factors included in our study are not diverse enough to adequately cover other cognitive domains, such as cognitive flexibility (switching) or language. We hope that the current work motivates future source-space M/EEG investigations of other cognitive domains. Another crucial aspect is that we investigated only a few hypothesis-driven connectome phenotypes (i.e., Transition Probability and Fractional Occupancy in specific bands) in the present study. These phenotypes were selected based on a preceding study, which stands as the sole investigation exploring the genetic impact on the time-varying characteristics of rapid connectome dynamics in source-space. However, it is likely that other dynamic features of the electrophysiological connectome may be individually specific and affect cognitive abilities. Future studies with sample sizes and analyses appropriate for more exploratory approaches may address a more extensive list of dynamic connectome features and behavioral measures.

Together, our findings establish that rapid, sub-second transitions between whole-brain connectome states, most notably the states’ sequencing in alpha, theta, and gamma bands, are associated with cognitive performance including visuospatial processing abilities. This evidence substantially extends previous associations of infra-slow dynamics and cognition (Vidaurre et al., 2017; Eichenbaum et al., 2020; Jun et al., 2022) to the rapid timescales at which most cognitive processes unfold. In light of the heritability of Transition Probabilities in resting-state electrophysiology (Jun et al., 2024; Submitted) our findings position these connectome features as potential endophenotypes for cognitive abilities. Such endophenotypes can delineate the neurobiological (specifically connectome-based) mechanisms via which genetics can shape cognitive abilities. Further, the identified connectome phenotypes may be of clinical relevance. Infra-slow (fMRI-derived) connectome dynamics are implicated in numerous psychiatric and neurological conditions (Preti et al., 2017) and explain cognitive abilities across different diagnostic categories (Zhu et al., 2021). However, the role of rapid electrophysiological connectome dynamics in cognitive and mental health is largely unexplored. Addressing their untapped potential, the identified phenotypes may be explored as connectome-based biomarkers of cognitive functioning and dysfunction with cost-efficient EEG.

## Acknowledgements

We thank Dr. Andre Altmann for his extensive guidance in analytic approaches, and Drs. Jonathan Wirsich and Thomas Alderson for guidance in data preprocessing. Computational resources for this work were provided by the Minnesota Supercomputing Institute at the University of Minnesota Informatics Institute. The Center for Magnetic Resonance Research (supported by Grant Nos. NIBIB P41 EB027061 and 1S10OD017974-01) at the University of Minnesota provided resources that contributed to the MRI-related results reported within this article. The original data collection of the data analyzed in this paper was funded by NIH grants R37 DA05147 and R01 DA036216. This work was partly supported by the National Institute for Mental Health (1R01MH116226 to Sepideh Sadaghiani).

