## Supplementary File for "Cognitive abilities are associated with rapid dynamics of electrophysiological connectome states"

**Supplementary Tables**

### Table S1. Cognitive Task Measures from Minnesota Twin Family Study

| **Cognitive Assessments** | **Task Description** | |
| --- | --- | --- |
|  | **Variable Names** | **Measured Performance** |
| WAIS-R Vocabulary and Block Design | Vocabulary Subtest: The examiner presents the test taker with a series of words, and the test taker is required to provide definitions for each word.  Block Design Subtest: The test taker is presented with a set of red-and-white blocks and a series of designs or patterns, progressively increasing in difficulty, and is then required to replicate the designs using the blocks. | |
|  | Verbal intelligence | The accuracy and depth of the word definitions. Verbal reasoning and vocabulary knowledge. |
|  | Nonverbal intelligence | Number of designs correctly replicated within the time limit. Non-verbal reasoning and spatial abilities. |
| WAIS-III Digit Symbol | Digit Symbol Coding: On the top of the page, a key with two rows (top: one-to-nine digits, bottom: a symbol for each digit) is presented. Test takers copy the symbols into spaces below a row of numbers according to the key.  Symbol Search: Two target symbols appearing on the left of a row are sought among an array of five symbols on the right. Test takers respond by either marking the identical symbol, or a “no” box (if the matching symbol is not present in the array). | |
|  | Processing Speed Index | Sum of subtest scores (Digit Symbol Coding and Digit Symbol Search Tasks). Ability to scan and process visuospatial stimuli and Attention. |
| Rey Auditory Verbal Learning Test | Trials 1 to 5: A sequence of 15 words (List A) is read aloud by an examiner, and the test taker is immediately asked to recall as many words on the list as possible.  List B: After that, a different set of 15 words (List B) is read to the test taker, who then is immediately asked to recall the words.  Trial 6: Immediately after List B, the test taker is asked to recall the words from List A.  Trial 7: Recall, unwarned, the words from List A after a delay of 30 minutes after Trial 6, during which time the test taker performs a series of nonverbal tasks. | |
|  | Total immediate recall | Sum of words immediately recalled after the List A is read to the test taker across Trial1 ~ Trial 5 |
|  | Short-delay recall | Trial 6 recall performance – Trial 5 recall performance |
|  | Long-delay recall | Trial 7 recall performance – Trial 5 recall performance |
|  | Learning over trial | Trial 5 – Trial 1 |
| Visuospatial Paired Associates Learning | Boxes are displayed on the screen and are “opened” in a randomized order. One or more of them will contain a pattern. The patterns are then displayed in the middle of the screen, one at a time and the test taker must select the box in which the pattern was originally located. | |
|  | Visuospatial Learning | Inversed value of total error numbers. Visuospatial learning and memory. |
| WAIS-III Digit Span | The digit sequences typically start with two digits and progressively increase in length.  Forward Digit Span: Test-takers repeat the read-out list of digits in the order.  Backward Digit Span: Test-takers repeat the read-out list of digits in reverse order. | |
|  | Forward Digit Span | Number of correct trials. Attention and Working memory. |
|  | Backward Digit Span | Number of correct trials. Working memory and Executive function. |
| Spatial Span Task | The stimuli sequences typically start with two squares with a variable order of colors and progressively increase in length.  Forward Spatial Span: Test-takers recall the sequence and location of a series of spatial stimuli in the order.  Backward Spatial Span: Test-takers recall the sequence and location of a series of spatial stimuli in reverse order. | |
|  | Forward Spatial Span | Number of correct trials. Attention and Working memory. |
|  | Backward Spatial Span | Number of correct trials. Working memory and Executive function. |
| Go/No-Go task | Test takers respond by pressing a button when they see a “go” signal and withhold a response when they see the “no-go” signal. The MCTFR version is a one-back go/no-go task: participants press a button to each stimulus in a sequence of two letters unless the current letter is identical to the letter on the immediately preceding trial. | |
|  | Prepotent motor inhibition | Inverse value of false alarm rate (%). Ability to inhibit prepotent motor response. |
| Iowa Gambling Task | Test takers are presented with four decks of cards labeled A, B, C, and D. Each deck contains a mix of gains and losses. Test takers are instructed to choose cards from these decks without knowing their characteristics to accumulate as much money as possible with feedback given after each selection. | |
|  | Decision under Uncertainty | Advantageous decision-making under uncertainty. Ability to learn from feedback and adjust their choices based on risk and reward outcomes. |
|  | Decision under Risk | Advantageous decision-making under risk. Ability to learn from feedback and adjust their choices based on risk and reward outcomes. |

WAIS-R; Wechsler Adult Intelligence Scale-Revised (WAIS-R).

**Supplementary Figures**

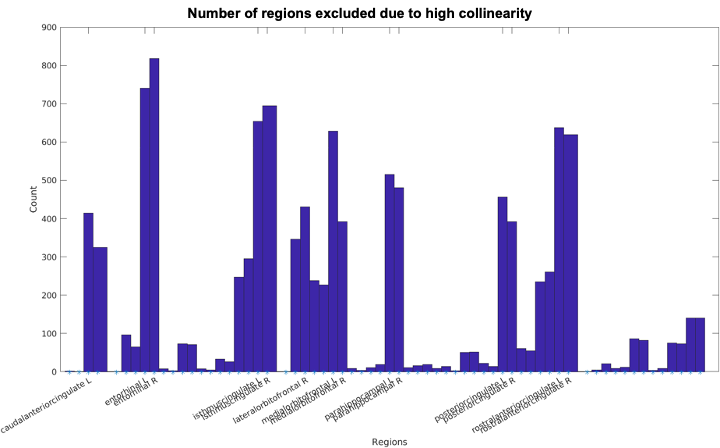

##### Figure S1. Top 14 regions with high collinearity.

To mitigate the source-leakage confound, caused by the blurring of point dipole sources and the spreading of signals across neighboring regions, we excluded regions, whose extracted signals were found to be highly collinear with others based on *qr* function in Matlab. Subsequently, 14 regions were excluded from the investigation: bilateral ‘*rostralanteriorcingulate’*, bilateral '*posteriorcingulate*', bilateral '*parahippocampal*', bilateral '*medialorbitofrontal'*, bilateral '*isthmuscingulate*', bilateral '*entorhinal*', '*lateralorbitofrontal R*', and '*caudalanteriorcingulate L'*.

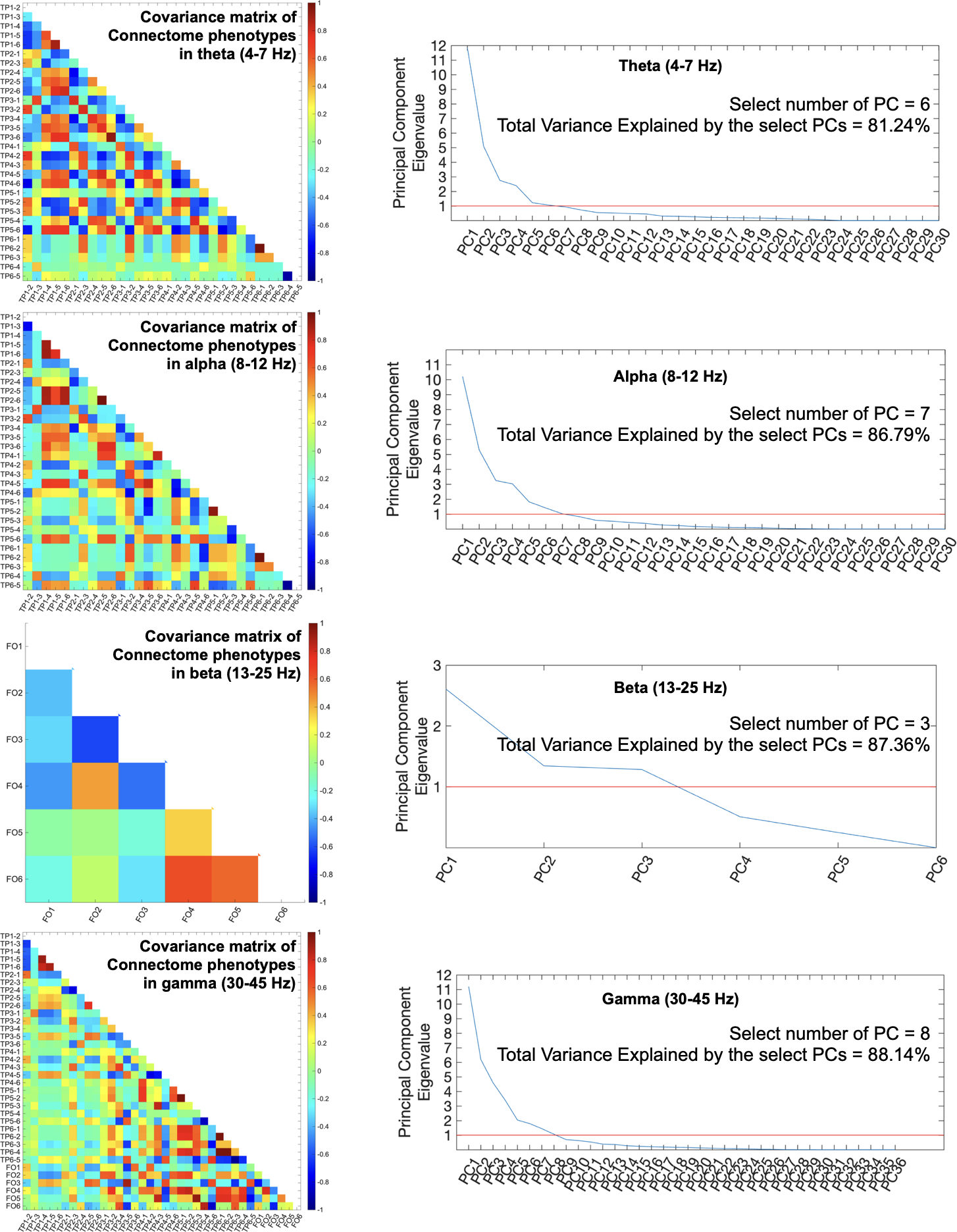

##### Figure S2. Overview of the principal component analysis on temporal features of the dynamic connectome.

(Left) Pearson’s correlation matrix for the temporal features of the dynamic connectome obtained from the six-state model, color-coded for Pearson’s correlation coefficient. (Right) Scree plot of the eigenvalues of principal components of the temporal connectome dynamics features for each frequency band. Red colored reference line (y = 1) indicates the cut-off value used for the present study.

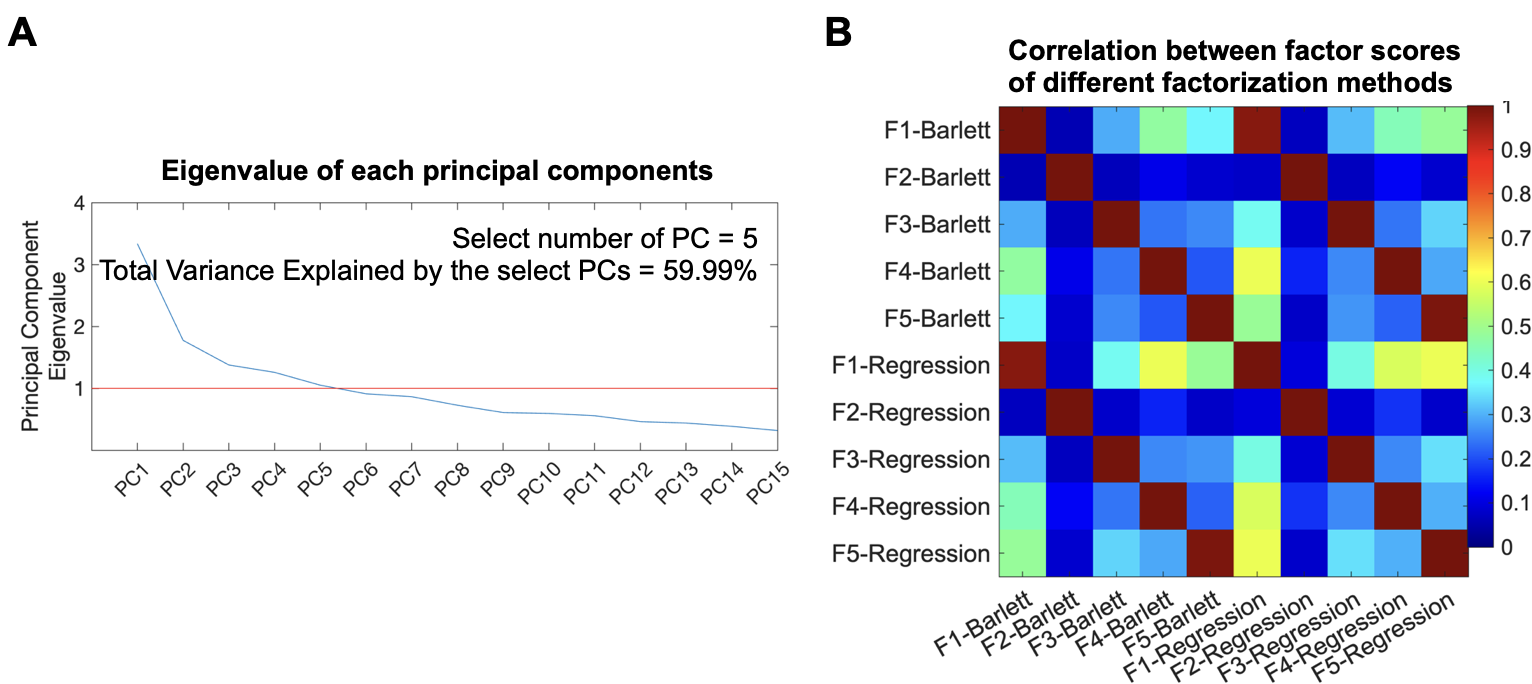

##### Figure S3. Overview of the cognitive dataset.

(A) Scree plot of the eigenvalues of principal components obtained from the covariance matrix of the cognitive measures, where red colored reference line (y = 1) indicates the cut-off value used for the present study. (B) Pearson’s correlation matrix for two sets of factor scores derived using Bartlett method (F1-Barlett to F5-Barlett) and ridge regression method (F1-Regression to F5-Regression). PC: principal component, F: Factor.

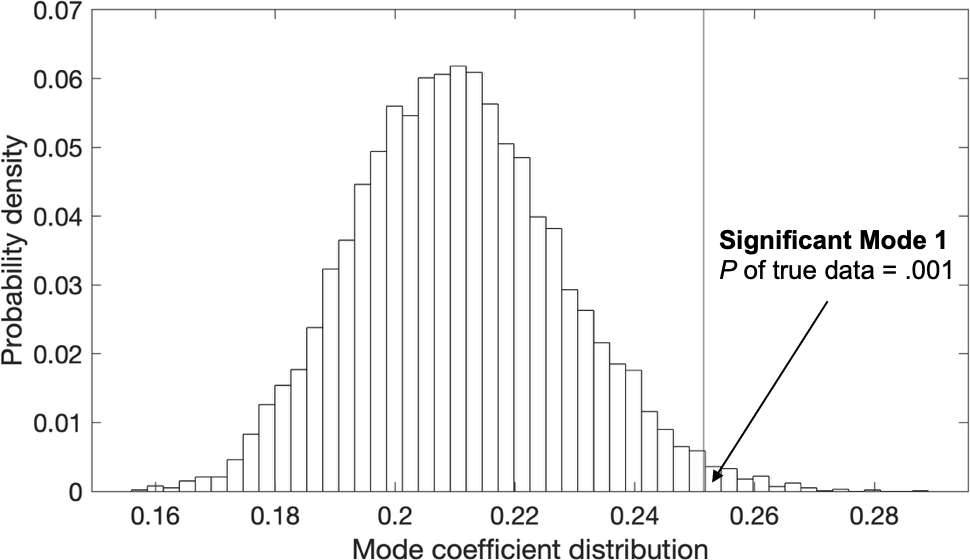

##### Figure S4. The distribution of random canonical variate pair correlation values

The histogram shows the distribution of random canonical variate pair correlation values from 10,000 permutations of the rows of U relative to V were performed, while maintaining the within-participant structure of the data. The mode of covariation identified in the real data, compared against the permutations, showed significance at *p* = .0012.
